# Fungal biodiversity in Arctic paleoecosystems assessed by metabarcoding of lake sedimentary ancient DNA

**DOI:** 10.1101/2021.11.02.462738

**Authors:** PA Seeber, B von Hippel, H Kauserud, U Löber, KR Stoof-Leichsenring, U Herzschuh, LS Epp

## Abstract

Fungi are crucial organisms in most ecosystems as they exert ecological key functions and are closely associated with land plants. Fungal community changes may therefore help reveal biodiversity changes in past ecosystems. Lake sediments contain DNA of organisms in the catchment area, which allows reconstructing past biodiversity by using metabarcoding of ancient sedimentary DNA. We developed a novel PCR primer combination for fungal metabarcoding targeting a short amplicon to account for length bias of amplification due to ancient DNA degradation. *In-silico* PCRs showed higher diversity using this primer combination than using previously established fungal metabarcoding primers. We analyzed existing data from sediment cores from four artic and one boreal lake in Siberia. These cores had been stored for 2–22 years and examined degradation effects of ancient DNA and storage time-related bias in fungal communities. Amplicon size differed between fungal divisions, however, we observed no significant effect of sample age on amplicon length and GC content, suggesting robust results. We also found no indication of post-coring fungal growth during storage distorting ancient fungal communities. Terrestrial soil fungi, including mycorrhizal fungi and saprotrophs, were predominant in all lakes, which supports the use of lake sedimentary ancient DNA for reconstructing terrestrial communities.

## Introduction

Fungi constitute the third-largest kingdom on Earth in terms of biomass, after plants and bacteria (Bar-On *et al*., 2018), and they have considerable effects on structure and functioning of most ecosystems. Fungi are essential to the survival, growth, and fitness of many organisms with which they form associations, including enumerable plant species, in almost all ecosystems (Brundrett, 2004; Finlay, 2008). Biodiversity of mycorrhizal fungi is known to affect plant community structures, thereby affecting entire ecosystems (van der Heijden *et al*., 1998; Clemmensen *et al*., 2013; Powell and Rillig, 2018). Particularly in environments which are notoriously nutrient-poor, e.g., the Arctic, plants are typically highly dependent on symbioses with mycorrhizal and endophytic fungi (Smith and Read, 2008). Apart from mycorrhizal symbioses, fungi exert various other key ecological functions in terrestrial and aquatic habitats, including decomposition of components of complex substrates such as cellulose and lignin, subsequent recycling of nutrients, and pathogenic effects on countless taxa of eukaryotes (Martin, 2014). However, compared to other kingdoms such as Animalia and Viridiplantae, knowledge on fungal diversity and distribution of fungal taxa and functional groups is relatively limited (Baldrian *et al*., 2021). The comparable lag in studies on fungi, particularly within their natural habitats, may be attributed, in part, to the microscopic dimensions of their vegetative bodies with hyphae of only a few microns in diameter, but also to the extreme taxonomic diversity within the fungal kingdom with a currently estimated 1–4 million species (Blackwell, 2011; Hawksworth and Lücking, 2017; Baldrian *et al*., 2021), compared to approximately 400,000 species of vascular plants. However, only approximately 120,000 species of fungi are thoroughly described and accepted (Hawksworth and Lücking, 2017).

One crucial problem in fungal systematics and taxonomy is that many fungi occur in morphs which differ drastically regarding their phenotypic appearance. Moreover, some fungal species can comprise multiple strains that can vary in their morphology, which has often led to their description as different species. Molecular approaches to assess biodiversity from environmental samples such as soils or sediments predominantly rely on metabarcoding, and can be used on modern or ancient DNA (reviewed by Ruppert *et al*., 2019). This method is generally a powerful tool for assessing species richness in an ecosystem (Deiner *et al*., 2017); however, metabarcoding of fungi is notoriously challenging due to the overwhelming taxonomic diversity of this group (Nilsson *et al*.; James *et al*., 2020; Lücking *et al*., 2020). Metabarcoding (Taberlet *et al*., 2012) depends on PCR amplification of a stretch of DNA with sufficient variation to allow for taxonomic assignment at the desired resolution but lies within two regions that are conserved enough to allow annealing of primers exclusively specific to (and only to) the respective taxonomic group. Regarding fungi, the most common metabarcoding regions are the two internal transcribed spacers (ITS) ITS-1 and ITS-2 of the ribosomal RNA genes (Schoch *et al*., 2012; Stielow *et al*., 2015). Even though resolution to species level based on ITS barcodes may be difficult, the current reference databases are most comprehensive regarding this marker, compared to others (Stielow *et al*., 2015; Lücking *et al*., 2020), and it is also used as the standard barcode by the International Barcode of Life consortium. The databases for the marker have increased tremendously in the past years, enabling more precise taxonomic assignments. Furthermore, the increase in available sequences potentially also offers the possibility to optimize metabarcoding assays in comparison to the primers that are currently in common use. Robust metabarcoding assays are of particular relevance for the analysis of potentially degraded environmental DNA, as recovered from ancient sedimentary deposits (Lydolph *et al*., 2005; Bellemain *et al*., 2013; Talas *et al*., 2021). Primers for this purpose have not been reevaluated in close to a decade (Bellemain *et al*., 2010, 2013; Epp *et al*., 2012), even though fungal metabarcoding may help reveal community turnovers and trace ecosystem changes on very long timescales (von Hippel *et al*., 2021).

Analyses of sedimentary deposits can reveal fungal community alterations and associated ecosystem changes over long periods of time. Investigating past fungal biodiversity changes by reconstructing paleoecosystems may generate insights regarding basal structural developments that are to date mostly overlooked in classical palynological approaches due to reliance on microscopic remains (Taylor and Osborn, 1996; Wood and Wilmshurst, 2013; Chepstow-Lusty *et al*., 2019). While molecular genetic methods are commonly used at present to investigate modern ecosystems (Heeger *et al*., 2018; Adamo *et al*., 2020), far fewer studies have been conducted on fungal biodiversity in paleoecosystems. Early studies concentrated on samples from permafrost soils (Lydolph *et al*., 2005; Bellemain *et al*., 2013), in which DNA preservation is optimal and which was in general an early target for sedimentary ancient DNA (Willerslev *et al*., 2003, 2014; Haile *et al*., 2009). DNA from these deposits showed a high potential for the analysis of past communities, but with certain peculiarities regarding fungi. In particular, Bellemain *et al*. (2013) suggested that the fungal DNA in permafrost potentially originated not only from ancient communities, but also from organisms that were still alive. Moreover, sample material (such as sediment cores) is frequently stored for long periods of time before DNA isolation and may thus be prone to growth of fungi (e.g., molds), which could distort ancient community signals. Thus, such storage effects must be considered.

Regarding ancient DNA of many other organismal groups, lake sediment cores have by now become the most commonly targeted environmental archive (Domaizon *et al*., 2017; Parducci *et al*., 2017). Lake sediments comprise organic and inorganic matter originating from the lake’s catchment (and beyond), including intracellular and extracellular DNA of a vast spectrum of organisms which has been shed into the environment and was subsequently translocated to the sediment by various physical processes. Under adequate conditions (*e.g*., low temperatures and neutral to slightly basic pH), integration into lake sediments may help preserve environmental DNA and shield it from degradation over considerable time spans (reviewed by Capo *et al*., 2021). Thus, sediment cores may contain a plethora of information with which past ecosystems and changes in biodiversity and community structures can be reconstructed (reviewed by Capo *et al*., 2021). So far, explicit fungal metabarcoding from lake sedimentary DNA has been performed in only one study, which used a multiplex PCR approach (Talas *et al*., 2021), and showed that this approach can be used to assess past fungal communities and processes in lakes and the surrounding terrestrial environment.

Here, we developed metabarcoding primers to investigate past fungal biodiversity, and we assessed the specificities of fungal sedimentary ancient DNA (sedaDNA) extracted from lake sediment cores in Siberia (for details see von Hippel *et al*., 2021). To this end, we re-evaluated the existing metabarcoding assays for their use in paleoecology and updated metabarcoding primers for used with sedimentary ancient DNA. Using this assay, we examined taxonomic resolution and richness as well as replicability in paleorecords of five Siberian lakes, four of which are located in the Arctic and one in boreal South-East Siberia. All cores reach back through the Holocene and beyond and were stored for different time periods (2–22 years) in storage facilities. With this set of cores we intend to assess potential biases due to sampling location, sample age and storage time before DNA extraction, and capture the common characteristics of fungal DNA from lake sediments across a vast geographical area. For one core from the Taymyr Peninsula, for which we have the best temporal resolution, we explored trends of diversity in greater detail.

## Results

### Primer design and evaluation

We evaluated potential combinations of newly designed and established metabarcoding primers *in silico* and three candidate primer pairs did not contravene any of the exclusion criteria (one pair in ITS-1 and two pairs in ITS-2; Tables 1 and Supplementary Table S1). Based on this, we chose a combination which produced short amplicons (mean length of 183 bp), showed high specificity to fungi (91% of the amplicons assigned to fungi), and amplified a high number of target sequences *in silico* (N = 383,992). This was the combination ITS67 (5’-ACC TGC GGA AGG ATC ATT-3’; this study) and 5.8S_Fungi (5’-CAA GAG ATC CGT TGT TGA AAG TT-3’; Epp *et al*., 2012). Most other primer combinations were excluded because they produced large numbers of off-target amplicons (mostly of plants) or amplicons which exceeded our chosen threshold of 250 bp (Table 1). For the chosen primers, *in silico* amplicons of Ascomycota (mean length 186 bp; N = 157,058) were shorter than those of Basidiomycota (mean 207 bp; N= 62,108), and fungi *incertae sedis* (including Mucoromycota) amplicons were, on average, 168 bp long (N = 14,308). The taxonomic resolution (i.e., the proportion of unambiguously identified taxa) of the amplified marker was 53% at species, 67% at genus, and 60% at family level, according to the ecotaxspecificity function of obitools software (Boyer *et al*., 2016). Regarding off-target amplification, the chosen primer combination produced 6,078 amplicons from the Viridiplantae database, and 386 amplicons from the Metazoa database.

**Table 1.**
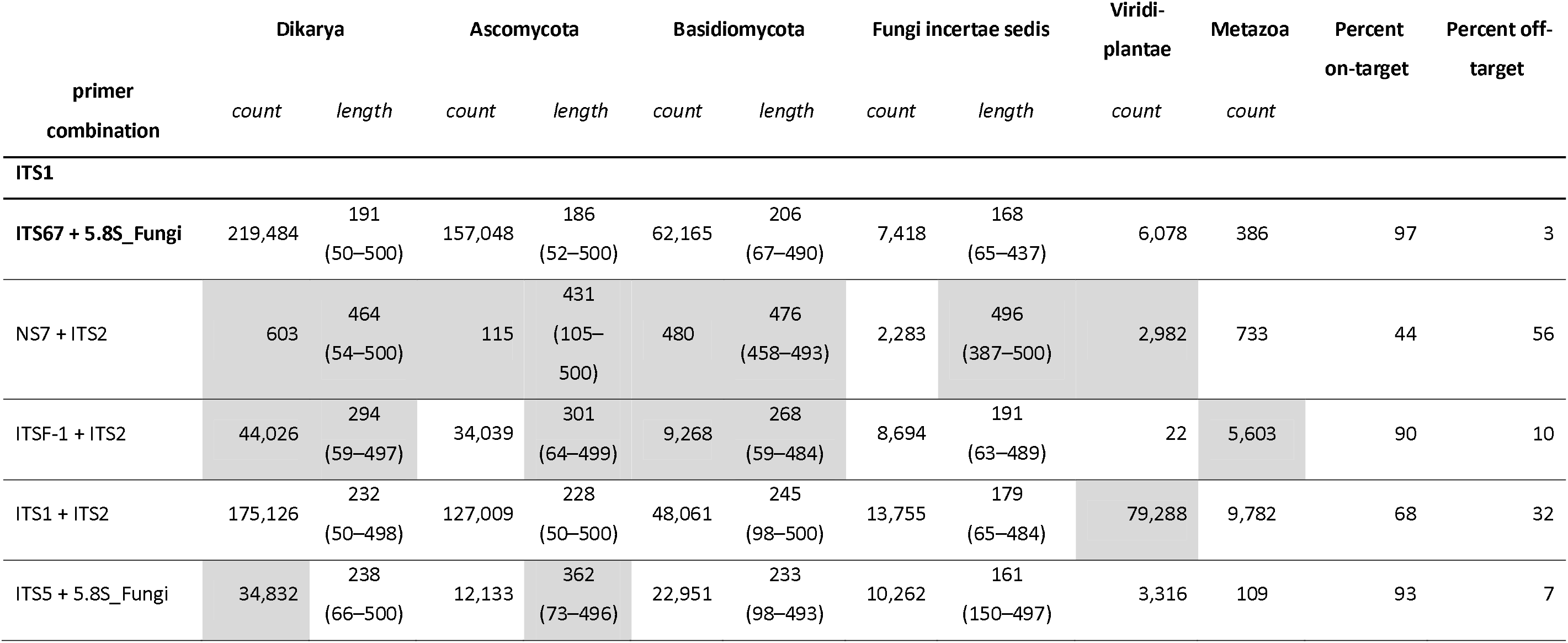
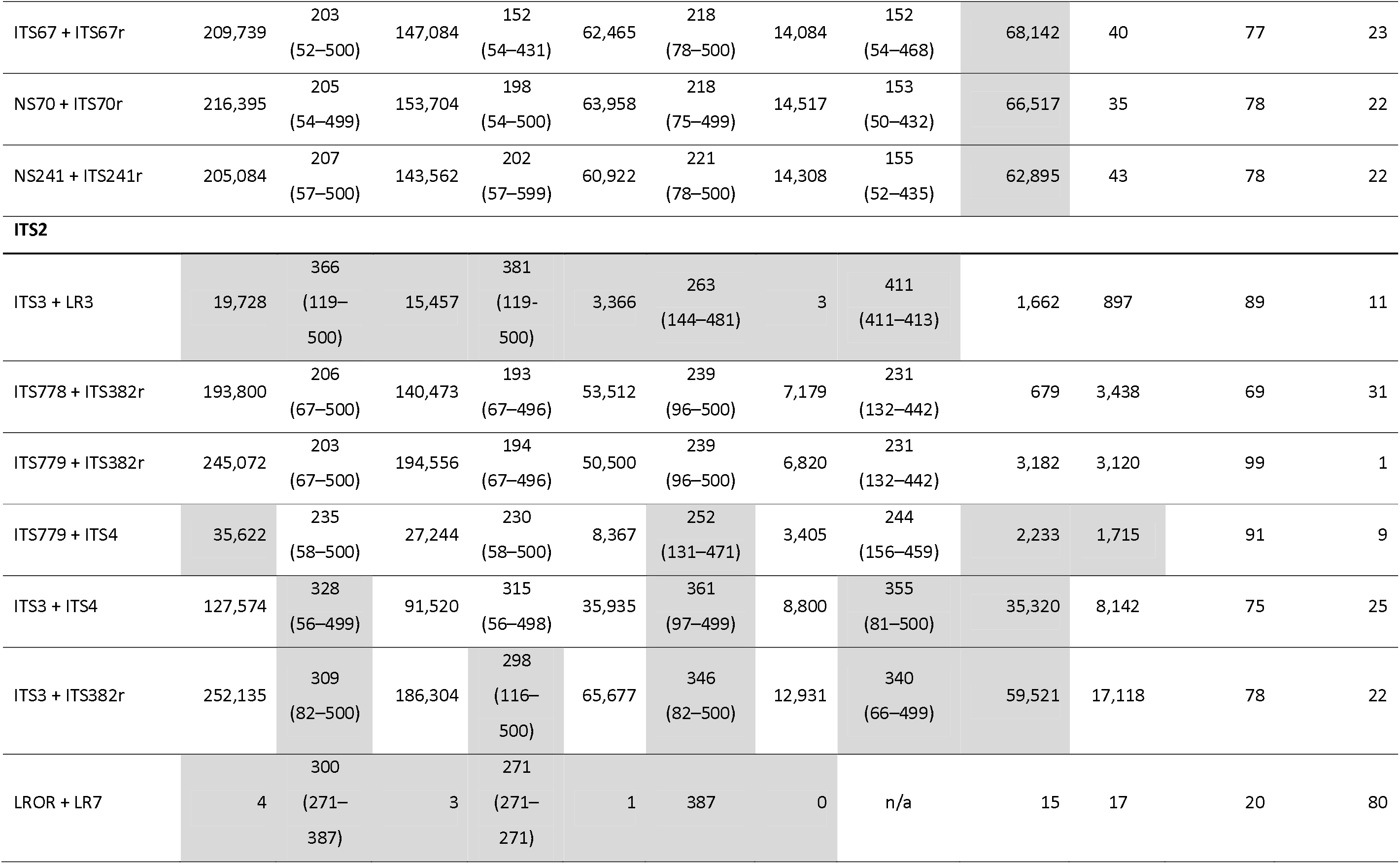
Evaluation of candidate primers for fungal metabarcoding using in silico PCRs. Targeted were the internal transcribed spacer (ITS)-1 (top section) and ITS2 (middle section), respectively, with EMBL release #142 as a reference database; shown are the counts of unique amplicons of the fungal subkingdom Dikarya, the divisions Ascomycota, Basidiomycota, and other divisions which are comprised on division rank by ‘Fungi incertae sedis’, as well as the off-target clades Viridiplantae and Metazoa, and average amplicon lengths (with the respective minimum and maximum, in parentheses; for fungal taxa). Exclusion of primer combinations for comprehensive fungal metabarcoding according to the specified criteria are indicated by gray shading. Bold print indicates the primers chosen for the metabarcoding experiments. Nucleotide sequences and references are shown in Supplementary Table S1.

Primers ITS67 and 5.8S_Fungi were tested *in vitro* in PCRs, and the reaction conditions were optimised using template DNA extracted from Museum vouchers of eight fungal taxa, including the geneera *Sordaria*, *Podospora*, *Cladnoia*, *Panaeolus*, *Coprinus*, *Coprinopsis*, and *Pilobolus*. All reactions produced amplicons in the expected size range.

### Evaluation of lake sediment core DNA for fungal paleoecology

#### Taxonomic resolution across the cores

After processing and filtering of the raw data including clustering at 97% (described in detail by von Hippel *et al*., 2021), the resulting 5,411 cluster centroids were subjected to taxonomic assignments with each database and to subsequent filtering, as indicated above. Across the 70 samples of five cores and using the UNITE database, 135 operational taxonomic unit (OTU) cluster centroids were retained (Supplementary Tables S2 and S3), which comprised 33 taxonomic orders (Table 2), 57 families, 79 genera, and 113 species. Regarding maximum taxonomic resolution, 121 (89%) OTUs were assigned to species level, 7 (5%) to genus level, 3 (3%) to family level, and 3 (3%) to order level. Using the EMBL database, 384 OTU cluster centroids were retained, comprising 51 orders, 100 families, 152 genera, and 188 species; 556 (50%) of the OTUs were assigned to species level, 275 (29%) to genus level, 66 (7%) to family level, and 13 (1%) to order level; 4% of the OTUs were assigned only to higher taxonomic levels. The overall weighted mean frequencies at class level were plotted across all samples of each lake (Fig. 1), and changes in taxonomic resolution (i.e., assignment to species, genus, family, class, and order level) over sample age are shown in Fig. 2.

**Fig. 1.**
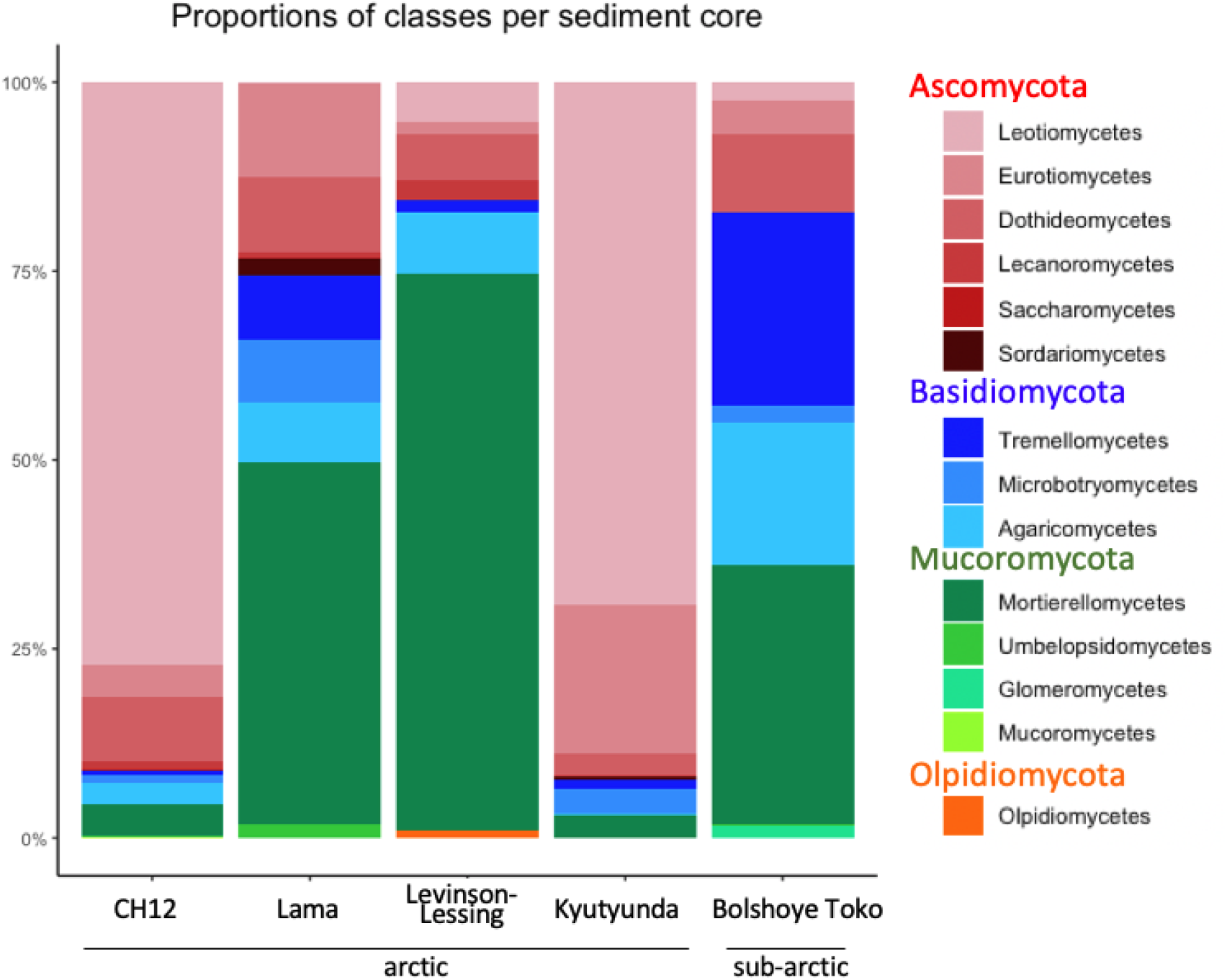
Proportions of fungal classes in lake sediments, averaged across all respective samples.

**Fig. 2.**
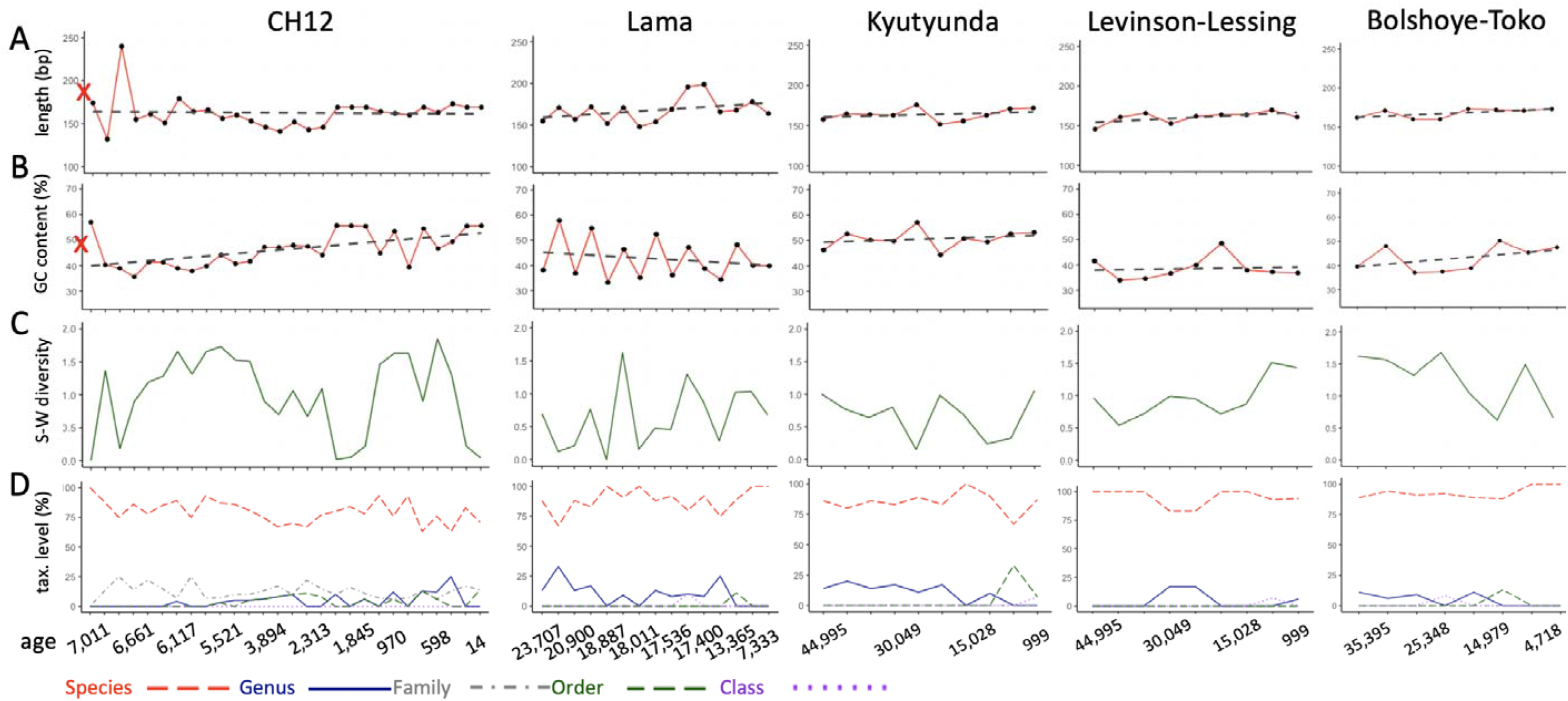
Effect of sample age on A) mean amplicon lengths, B) weighted mean GC content, C) Shannon-Wiener diversity indices (DI) of fung al taxa on order level, a D) taxonomic taxonomic resolution, i.e., the percentage of OTUs assigned on species (red), genus (blue), family (gray), order (green), and class (purple) level. R crosses on the y-axes of panels A and B indicate the respective values of the in silico PCR output (183 bp length and 74% GC content).

**Table 2.**
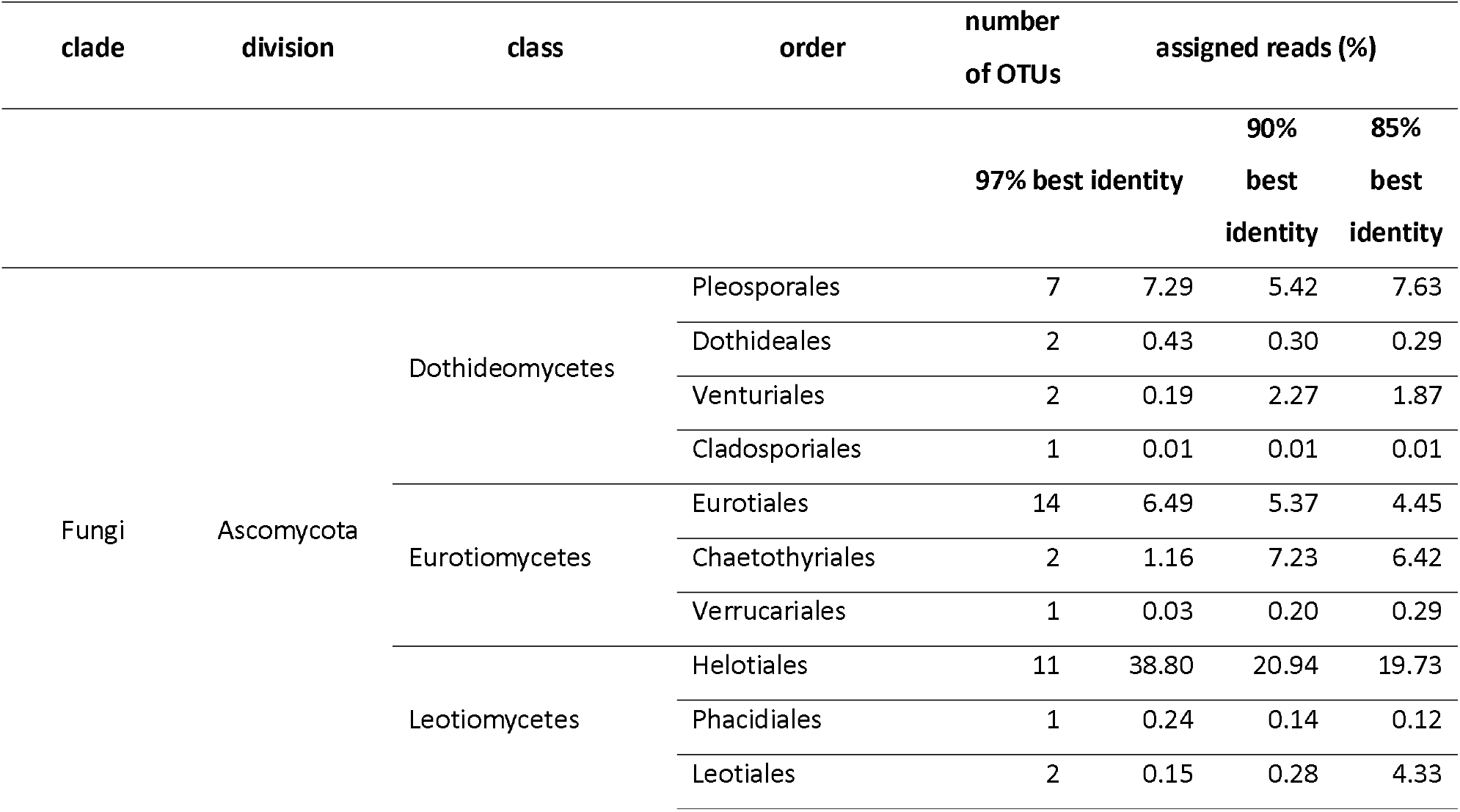
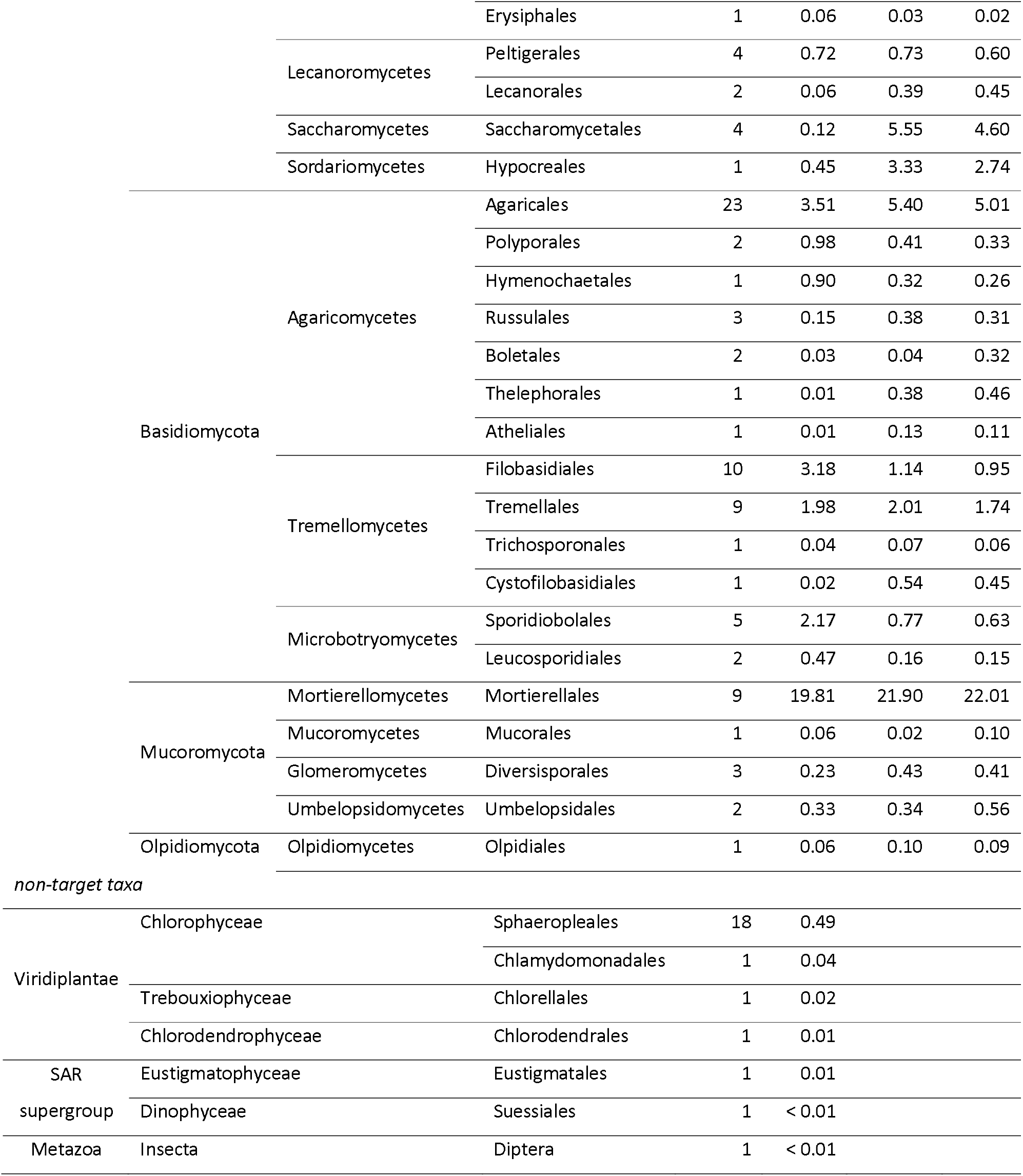
Numbers of OTUs and percent of respective reads of fungi (UNITE database) and non-fungi taxa (EMBL database; lower section) per taxonomic order. Percentages are based on the sum of assigned reads in the respective database.

#### Comprehensiveness: rarefaction and accumulation curves

Rarefaction analysis was performed to estimate OTU richness as a function of sampling effort, i.e., sequencing depth, based on the minimum number of observed sequence counts. Most rarefaction curves showed an asymptotic course, suggesting that sequencing depth was sufficient (Supplementary Fig. S1). To test whether the number of PCR replicates was sufficient to assess taxonomic richness per sample, we produced OTU accumulation curves of the core with the highest temporal resolution (from lake CH12; [Klemm *et al*., 2016]). These curves did not show saturation over the maximum of six replicates (Supplementary Fig. S2).

#### Amplicon length and GC content to assess bias through degradation

Average amplicon length as a function of age per core was tested to assess the effect of age on DNA degradation. This suggested only negligible decreases in mean amplicon length over time (Fig. 2A); in none of the five cores did the linear regression indicate a significant effect of age on amplicon length, regardless of taxonomic assignment. Degradation processes linked to the age of DNA also lead to an increase in GC content (Dabney *et al*., 2013); in the current study, amplicon GC content ranged from 10% to 63% (mean 43% ± 10%), and we observed no bias towards higher GC content in older samples. In core CH12, GC content rather decreased significantly with age (est. −0.19; t = −3.33; p = 0.003), while no significant effect of age on GC content was observed in cores of the other four lakes (Fig. 2B). By comparison, the GC content in the *in silico*-PCR output was 47%.

#### General taxonomic composition of fungi in Siberian lake sediment cores

The predominant taxa (Table 2 and Fig. 1) in all lakes were terrestrial saprotrophs, mycorrhizal fungi and other soil fungi, e.g., OTUs assigned to the genera *Mortierella* (order Mortierellales) and *Inocybe* (Agaricales) (Domsch *et al*., 2007; Varma *et al*., 2017), respectively. We found very few sequences of fungi that were determined to be aquatic, i.e. OTUs assigned to *Alastospora* sp. (Ascomycota). These were restricted to four samples of Lake Levinson Lessing and comprised less than 4000 reads in the total dataset.

The dataset also contained reads of off-target taxa (i.e., non-fungi) which were assigned using the EMBL database. Out of 936 EMBL-assigned OTUs, 24 OTUs at a total of 59,902 reads were assigned to non-fungi taxa, which belonged to the clades Viridiplantae (21 OTUs; 55,777 reads), the SAR supergroup (2 OTUs; 2,788 reads), and Metazoa (1 OTU; 1,337 reads; Table 2).

Reads produced from extraction blanks and non-template PCR controls were processed as described above and were assigned using the UNITE databases, which showed assignment of 141,746 reads to 34 OTUs. Three of the five most abundant taxa in the PCR and extraction controls (*Wickerhamomyces*, *Candida*, and *Pichia*) did not occur at all among assigned sample reads (Supplementary Table S2), and only the genera *Aspergillus* and *Gryganskiella* occurred in total numbers of > 10.000 reads in extraction blanks and non-template controls.

### Diversity of fungal paleo-communities from lake CH12

For core CH12, for which we have the highest temporal resolution, and which has previously been analyzed for pollen and plant DNA (Klemm *et al*. 2016, Epp *et al*. 2018), we conducted a more indepth analyses of diversity. We examined Bray-Curtis dissimilarities between PCR replicates of all samples of core CH12 to test whether within-sample variation is smaller than between-sample variation. The MRPP analysis suggested significantly lower dissimilarity between PCR replicates of one sample (64.97%) than between those of different samples (88.46%; p = 0.001; within-group agreement = 0.26; observed delta 0.6503; expected delta 0.8774). In the graphical cluster visualization based on Bray-Curtis distances, however, most replicates clustered together regardless of samples, likely due to the predominance of specific taxa in most samples (not shown). We therefore repeated this analysis after excluding a) the most abundant taxa (> 50.000 reads in total) or b) the least abundant taxa (< 50.000 reads in total); however, this analysis did also not visually differentiate samples based on similarities of the PCR replicates.

Across all cores, the fungal divisions Ascomycota, Basidiomycota, Mucoromycota, and Olpidiomycota were represented and accounted for approximately 62%, 15%, and 23%, and 0.06% of the fungal reads, respectively (Table 2). Amplicons of Basidiomycota were significantly longer (179 ± 44 bp) than those of Mucoromycota (153 ± 24 bp; ANOVA F = 3.383; p = 0.037; Tukey’s test p = 0.04). Amplicon lengths of Ascomycota (169 ± 22 bp) did not differ significantly from that of Basidiomycota and Mucoromycota. The single OTU assigned to Olpidiomycota was 104 bp long. In the case of CH12, we found that Ascomycota were more abundant in younger samples and that the proportion of Mucoromycota (shortest amplicons) was higher in older samples; however, Basidiomycota (longest amplicons) also appeared to be more abundant in older samples (Fig. 3). Shannon-Wiener diversity indices showed considerable variation in fungal communities over time (Figs. 2C). In core CH12, OTUs were assigned to a total of 23 orders, and taxonomic richness ranged from 1 (in the oldest sample, dated 7,011 years) to 14 (sample 5,521 years; mean 8 ± 3) with particularly low diversity in samples dated 1,976, 1,845, and 14 years (diversity indices 0.02, 0,05, and 0.03, respectively) even though the respective OTU’s were assigned to 7, 9, and 6 orders; this discrepancy between richness and diversity is due to the dominance of OTUs assigned to the order Helotiales in these samples (> 99%, each).

**Fig. 3.**
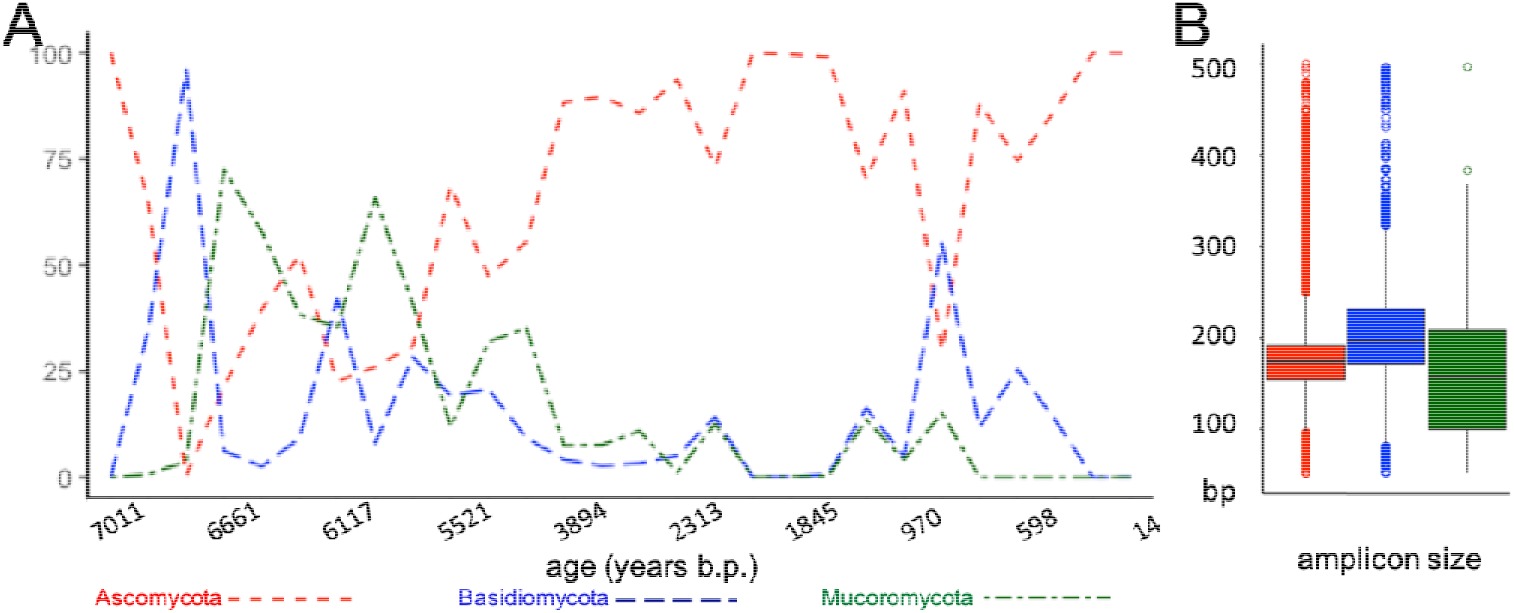
Changes in proportions of the three major divisions Ascomycota (red), Basidiomycota (blue), and Mucoromycota (green) over sample age (A), and respective fragment lengths (B); bBoxes indicate 2nd and 3rd quartiles, center lines indicate median values, upper (and lower) whiskers extend to the highest (and lowest) value within 1.5-times the inter-quartile range. Data points beyond the end of the whiskers are shown as open dots.

We found no evidence of introduced bias in fugal communities due to growth of molds post-coring. The cumulative overall proportions of the typical mold genera *Aspergillus*, *Cladosporium*, *Mucor*, and *Penicillium* in the cores were as follows: 1.37% in the Levinson-Lessing core (2017), 4.21% in the Bolshoye Toko core (2013), 3.72% in the CH12 core (2011), 15.66% in the Kyutyunda core (2010), and 5.76% in the Lama core (1997). The respective proportions remained relatively stable over time and did not appear to increase in deeper layers (Supplementary Fig. S3). By comparison, the proportion of these genera the *in silico* PCR output was 3.87%.

## Discussion

### Preservation biases and potential contamination

Our results confirm that ancient DNA from lake sediment cores is a reliable archive of fungal paleocommunities, and these can be reconstructed from boreal to arctic sites with ages up to 44,995 years. Fungal paleo-communities from lake sediments have also been recently reported from temperate conditions and spanning the Holocene (Talas *et al*., 2021). Our putatively authentic ancient fungal DNA, attributed primarily to taxa from the terrestrial surroundings of the lake, was in part recovered from cores that had been stored in a storage facility under non-frozen conditions for multiple years prior to sampling for DNA, underlining the value of sediment core collections also for work with sedimentary ancient DNA. For the ITS1-amplicon targeted by our primers, the average length of Ascomycota was shorter than that of Basidiomycota, both *in silico* using ecoPCR on the EMBL database and in the sequencing results (198 vs. 219 bp and 169 vs 179 bp, respectively), which is in line with the results of a previous study (Bellemain *et al*., 2010). This difference was smaller in sequencing reads from sedaDNA than in *in silico* PCR products, which could possibly be attributed to fragmentation of DNA in the environment. However, our results suggest only minor, non-significant declines in amplicon lengths over time in each sediment core (Fig. 2), indicating that the results are not impacted by modifications of amplicon lengths.

A further source of bias linked to degradation can potentially change GC content of the DNA. GC content can increase with age of DNA due to degradation impacting primarily GC-poor fragments (Dabney *et al*., 2013). In the current study, GC content ranged from 10% to 63%, and we observed no bias towards higher GC content in older samples. Contrary to our expectations, we found a decrease in GC content with age in one lake (CH12). This may, however, be an incidental result considering that no significant effect of age on GC content was observed in cores of the other four lakes (Fig. 2B). Taken together, the lack of GC bias corresponding to sample age suggests that degradation of arctic and subarctic sedaDNA may not be problematic for fungal metabarcoding. As we worked with cores that were not sampled for DNA soon after coring, but had in part been stored for 2–22 years prior to collecting and freezing DNA samples, we furthermore screened our results for the occurrence of taxa that point to recent fungal growth in or on the cores (e.g., dominance of molds). Growth of molds during storage of the cores or within the sediment would be expected to increase the proportion of these taxa, however, we found little indication for such a bias in our samples (Supplementary Fig. S3). Even though the second-oldest core (lake Kyutuynda; from 2010) showed the highest cumulative proportion of the four mold genera *Aspergillus*, *Cladosporium*, *Mucor*, and *Penicillium* (15.66%), this was only due to two samples with particularly high abundance, whereas in the core from 1997 (Lama), a substantially lower proportion of these taxa was observed (5.76%). Thus, long storage times of sediment cores are not necessarily problematic for reproducing ancient fungal communities.

### Characteristics of the optimized sedaDNA ITS1 metabarcoding assay

A large part of the ancient DNA pool in sediment cores is typically reduced to fragments well below 50 bp (Pedersen *et al*., 2016; Parducci *et al*., 2017; Lammers *et al*., 2021) and will not be traceable at all through PCR, but when employing a metabarcoding approach, short amplicons are preferable. We therefore aimed for metabarcoding primers to amplify the shortest possible amplicons, so as to account for potential length bias due to fragmentation of DNA with age, while providing a high specificity to fungi, as well as a high taxonomic resolution. Diversity estimates of fungal communities (also including lichenized fungi) based on metabarcoding may be considerably biased by the choice of the barcode locus (Tedersoo *et al*., 2015; Tedersoo and Lindahl, 2016; Banchi *et al*., 2018), and ITS-2 has been suggested to be preferable over ITS-1, particularly with respect to taxonomic assignment of lichenized fungi and of Basidiomycetes (Tedersoo *et al*., 2015; Banchi *et al*., 2018). Thus, a combined approach using both ITS loci would be most reliable. However, the greater fragment length of ITS-2 may introduce amplification bias when ancient DNA is concerned. Our results using the primer combination optimized *in silico* suggested high taxonomic diversity at relatively short amplicon lengths (mean length of 183 bp), without discernible introduction of biases against one division. Due to the structure of the ITS region and its strong length variation, we could not design a universal primer pair for all fungi with a shorter amplicon, but our results suggest no bias in results due to group-specific amplicon length differences. Compared to an assay used recently for fungal DNA from a lake sediment core (Talas *et al*. 2021), the amplicon we propose appears to be substantially shorter than the fragment amplified by *in silico* PCR using the ITS3-ITS4 primer combination (mean length 338 bp; Table 1). In the study of Talas *et al*. (2021), greater length did not seem to compromise the results, but this may not universally be the case, and a different study recovered largely similar communities using ITS-1 and ITS-2 markers (Blaalid *et al*., 2013).

Despite its short length, the taxonomic resolution offered by our metabarcoding fragment was high not only *in silico*, but also in the sedaDNA results, with a high number of OTUs assigned to species or genus level. The exact number of assigned OTUs, and the taxonomic level reached, was highly dependent on the database used, underlining the importance of reference sequence collections. In general, and particularly for a taxonomic group as highly diverse as fungi, limited availability of reference sequences is one of the most crucial factors curbing the potential of metabarcoding communities from bulk samples. Moreover, resolving taxonomic diversity in bulk environmental samples may also be confounded due to continuously developing taxonomies and phylogenies and subsequent disagreement between databases regarding taxonomic levels and assignments; e.g., the EMBL database still uses the former division Zygomycota which as per the current standard was split into the phyla Mucoromycota and Zoopagomycota (Spatafora *et al*., 2016), as implemented in the UNITE database.

Despite these persistent, inherent challenges to fungal metabarcoding, the observed diversity regarding proportions of fungal divisions resembles the generally known proportions. The numbers of described species within the Ascomycota and Basidiomycota as the two largest divisions of the fungi kingdom exceed 60,000 and 30,000, respectively, whereas all other divisions comprise fewer than 2,000 known species, each (Naranjo-Ortiz and Gabaldón, 2019); we identified 55 OTUs of Ascomycota, 61 of Basidiomycota, and 15 of Mucoromycota, suggesting that the primer pair shows good universality to capture fungal diversity. We also screened the *in silico* PCR results for specific groups that have previously been targeted, or have been prominent, in sediment core studies. These are aquatic taxa, and in particular Chytridiomycota, which constituted a major component in the recently published study of Talas *et al*. (2021), as well as coprophilous fungi, such as *Sporormiella* and *Sordaria*, which are used in palynological studies as proxies for the presence of mammals (Gill *et al*. 2009, 2013, Davies 2019). These taxa occurred in the output of the *in silico* PCR and should thus potentially be retrieved by our assay.

### Potential of lake sediment fungal DNA for paleoecology

Fungi which are specific regarding their associations with plants or regarding other environmental conditions such as temperature may be indicative of local plant communities and climate, respectively. Based on the ecological conditions in the environment of the sampled lakes and on previous morphological and molecular studies in various ecosystems, we expected fungal taxa belonging to different ecological functional groups such as terrestrial saprotrophs, mycorrhiza fungi, coprophilous fungi, and aquatic taxa (Booth, 2011; Grau *et al*., 2017; Botnen *et al*., 2020) which may indicate ecosystem structures and environmental changes. Some examples of such ecosystem indicators are provided here, however, ecological interpretations of these data are made elsewhere (von Hippel *et al*., 2021).

The main fungal divisions were Ascomycota, Basidiomycota, and Mucoromycota, and we found relatively few reads (0.26%) of taxa which may be assigned to Glomeromycota, according to previous taxonomic systems, which is in line with the results of a recent metabarcoding study on fungal diversity from sediment of a lake in Eastern Latvia at 56.76 N, 27.15 E (Talas *et al*., 2021) with respect to terrestrial taxa. The overall composition differed between cores, which suggests local effects on past communities in the catchment of the respective lake; however, each of them seemed to reflect terrestrial communities. The fifteen most abundant taxa in each lake comprised terrestrial saprotrophs and mycorrhizal fungi (e.g., the genera *Mortierella* and *Inocybe*, respectively), which supports the use of lake sediment for reconstructing terrestrial fungal communities. OTUs assigned to the genus *Mortierella* were among the fifteen most abundant genera in each core, and these fungi typically occur as saprotrophs in soil, on decaying leaves, and other organic material (Domsch *et al*., 2007). Among mycorrhizal fungi, the genus *Inocybe* (Deacon *et al*., 1983) dominated some of the samples of lake CH12 and also occurred sporadically in samples of other lakes (von Hippel *et al*., 2021).

Contrary to our expectations, and in contrast to one of the main uses of fungal remains in paleoecology using classical, microscopic techniques, we observed no taxa which may be considered obligate coprophilous fungi (e.g., *Sporormiella* sp., *Preussia* sp., and *Sordaria* sp.). Spores of these taxa are used in morphological studies as proxies for herbivore presence (Davis and Shafer, 2006; Raper and Bush, 2009; Feranec *et al*., 2011). These taxa were recovered previously by Bellemain *et al*. (2013), using fungal metabarcoding on ancient permafrost, however, these deposits were of purely terrestrial origin. A high proportion of samples from one of the analyzed sites (Main River) also yielded mammal DNA (Willerslev *et al*. 2014), indicating a particularly high abundance of megafauna at this site. However, our data do not support the use of fungal DNA from lake sediment cores as a proxy for the presence and abundance of mammals in a similar way to microscopic analyses of spores. Future comparative morphological and molecular investigation may help elucidate the discrepancy between the approaches. In terms of lichens, which constitute an important part of Arctic vegetation and are crucial as sustenance of herbivores (Kumpula, 2001), we found some 16,000 reads assigned to *Peltigera* sp., which occurred only in cores of Arctic lakes. Other common lichen taxa such as *Cladonia* sp. or *Stereocaulon* sp. were not observed, which may be due to absence of DNA of these taxa in the sediment cores, absence of these taxa in the respective ecosystems, or biases inherent to ITS-1-based metabarcoding of fungi (Tedersoo *et al*., 2015; Banchi *et al*., 2018).

While the lack of coprophilous taxa in the molecular lake sediment records may be viewed as a drawback compared to permafrost deposits, their overall use in paleoecology seems more straightforward as they seem to be less confounded by potentially living organisms. Permafrost deposits contained a relatively high proportion of psychrophilic taxa (Bellemain *et al*. 2013), which are not necessarily ancient and which give no further paleoecological information in an area of cold ground temperatures. In our set of lakes, we found two OTUs assigned to the genus *Leucosporidium* which may be considered psychrotolerant. These were among the most abundant taxa in several samples of lake CH12, and showed considerable variation through the core (0%–12% of the respective read proportions; mean 3% ± 4%). Here, these sequences thus may be indicative of climatic changes and show a potential value as paleoenvironmental proxy.

In comparison to the recently published dataset on the temperate lake from Eastern Latvia (Talas *et al*. 2021), one striking difference is that we observed relatively few (predominantly) aquatic taxa, e.g., *Alatospora* sp. (3,227 reads in 4 samples). This could reflect the actual situation in the Arctic and Subarctic lakes target here, as little is known on aquatic fungi in this part of the world. A previous study discovered a substantial amount of aquatic fungi in Scandinavian lakes (Khomich *et al*., 2017), however, it is also possible that the aquatic taxa that do occur in the areas of the current study are underrepresented in the current reference databases, and their sequences were therefore not assigned.

This points to an aspect that cannot be fully resolved at this point: comparing taxonomic diversity and relative abundance between studies is not straight-forward due to a) differences between primers (and inevitable PCR bias), b) discrepancies between analysis pathways (including data filtration assignment algorithms, and comprehensiveness of reference databases), and c) differences between ecosystems and sampling substrates (e.g., Arctic versus non-Arctic systems, and lake sediments versus permafrost or soil cores).

Taken together, however, our results support the use of sedimentary ancient DNA from lake sediments for reconstructing past fungal communities. The observed community changes may hold valuable information on ecosystem changes regarding abundance of host plants of mycorrhizal or pathogenic fungi, climatic changes, and other ecological functions exerted by specific functional groups of fungi (for details, see von Hippel *et al*., 2021). Ecological interpretations, however, should be made with caution due to currently limited knowledge on the ecology and ecosystem functions of numerous fungal taxa. Further research is thus needed to link alterations in past and recent fungal communities and with changes on the scale of ecosystems, which may require further elucidating the ecology of crucial fungal taxa.

### Experimental procedures

#### Primer design and evaluation

Fungal ITS databases increased tremendously in the past years; for example, the established fungi primer pair ITS1F/ITS2 (Gardes and Bruns, 1993) produced 4,658 sequences from EMBL release #102 (Epp *et al*., 2012) but 154,168 sequences from EMBL release #138, with the same settings. Moreover, primers which are currently in use have not been reevaluated in close to a decade (Epp *et al*., 2012; Bellemain *et al*., 2013). We thus designed novel PCR primers for the ITS-1 and ITS-2 regions and tested these *in silico* together with established primers according to Bellemain *et al*. 2013 and Epp *et al*., 2012. Briefly, six databases were created by running an *in silico* PCR on the complete EMBL release #138 using ecoPCR software (Ficetola *et al*., 2010) and with the fungi barcoding primers of Epp *et al*. (2012). We allowed for three mismatches except for the last two nucleotide (nt) positions at the 3’-end of each primer and limited product size to 50–2,000 nt, apart from one amplicon database which spanned the entire ITS-1–5.8S–ITS-2-region and was permitted a maximum product size of 4,000 nt (primers ITS1 and ITS4). On each of these six amplicon databases, we used ecoPrimers software (Riaz *et al*., 2011) to identify potential PCR primers which had to exactly match 70% of the target sequences and match 90% of the target sequences with a maximum of 3 mismatches; product size was limited to 50–500 nt. From the output of each database, we selected primer pairs with a maximum difference in melting temperatures of 4 °C. These primers were then tested in terms of universality within the largest fungal divisions, *i.e*., Ascomycota, Basidiomycota, and other fungal taxa collated as *‘incertae sedis’* using in silico PCRs; the produced amplicons were dereplicated using the command ‘obiuniq’ of the obitools software (Boyer *et al*., 2016). Primers that showed a strong bias against any of these three divisions were eliminated. To exclude amplification of non-target sequences, we tested the candidate primers on databases containing only Viridiplantae and Metazoa, respectively; primers that produced a large number of products from these databases (*i.e*., more non-target sequences than sequences from any of the three fungi divisions) were eliminated. A total of 15 primer pairs (8 targeting ITS-1 and 7 targeting ITS-2) was tested in silico (Table 1), and primer combinations were excluded (1) if the mean amplicon size exceeded 250 bp, (2) if the subkingdom Dikarya as the largest phylum was represented by fewer than 50,000 unique amplicons, (3) if Basidiomycota and Fungi *incertae sedis* amplicons were more abundant than those of the largest division Ascomycota, and (4) if the number of off-target taxa (Viridiplantae and Metazoa) exceeded 10% of the number of Dikarya amplicons.

The following primer combinations targeting ITS-1 and ITS-2, respectively were considered most promising: ITS67 (this study) and 5.8S_fungi (Epp *et al*., 2012) as well as ITS779 and ITS382r (this study). For the metabarcoding experiments, we selected the combination ITS67 (this study) and 5.8S_fungi as this produced slightly shorter amplicons than the other combination and appeared to produce a somewhat more even ratio of Ascomycetes and other divisions.

Primers ITS67 and 5.8S_Fungi were tested *in-vitro* using DNA isolated from museum vouchers. Samples of the following species was obtained from the Natural History Museum of Oslo, Norway – *Sordaria aclina*, *Podospora fimicola*, *Podospora pyriformis*, *Cladnoia rangiferina*, *Panaeolus fimicola*, *Coprinus sterquilinus*, *Coprinopsis cinerea*, and *Pilobolus chrystallinus*. DNA was isolated from these specimens using the NucleoSpin Plant II kit (Macherey Nagel, Düren, Germany) according to the manufacturer’s instructions. The PCR reaction mix included 0.25 μL Platinum Taq DNA-Polymerase High Fidelity (Thermo Fisher Scientific; 5 U/μL), 12.75 μL ultrapure water, 2.5 μL 10 × buffer, 2.5 μL dNTPs (2.5 mM), 1 μL BSA (20 mg/mL; New England Biolabs, Frankfurt, Germany), and 1 μL MgSO_4_ (50 mM), and 1 μL of each PCR primer (5 μM). The following thermocycling protocol was used: 94 °C for 5 min, followed by 40 cycles of 94 °C for 20 s, 56 °C for 20 s, and 68 °C for 30 s and final extension at 72 °C for 10 min.

### Evaluation of lake sediment core DNA for analyses of fungal paleoecology

#### Sampling and DNA extraction

The laboratory works for this data set were performed at the Palaeogenetic Laboratory at AWI in Potsdam (von Hippel *et al*., 2021). We used sedaDNA isolated from sediment cores of five lakes in the Russian Federation, four of which are situated in the Arctic: a) a lake termed CH12 located in the Siberian Taymyr region (Khatanga, Russian Federation; 72.399° N, 102.289° E; 60 m a.s.l., collected in 2011; chronology of the core described previously [Klemm *et al*., 2016]; ages are shown as years before present [b.p.]), b) Lake Lama, Taymyr peninsula, 69.520° N, 89.948° E; 53 m a.s.l., collected in 1997, c) Lake Levinson-Lessing (74.512° N, 98.591° E; 47 m a.s.l.), Taymyr peninsula, collected in 2017, d) Lake Kyutyunda (69.630° N, 123.649° E; 66 m a.s.l.), Yakutia, north-eastern Siberia, collected in 2010, and e) Lake Bolshoye Toko (56.265° N, 130.530° E, 903 m a.s.l.) south-eastern Siberia (Neryungrinsky District, Sakha Republic), collected in 2013. For details on the cores and age-depth models, see von Hippel *et al*. (2021).

DNA was isolated from approximately 2–5 g sediment using the PowerMax Soil DNA Isolation kit (Qiagen, Hilden, Germany), and DNA extracts were purified and normalized to 3 ng/μL using a GeneJET Genomic DNA Purification Kit (Thermo Fisher Scientific, Bremen, Germany); for details see von Hippel *et al*. (2021). A total of 70 samples was used (CH12 core: 28 samples; Lake Lama core: 15 samples; Lake Levinson-Lessing core: 9 samples; Lake Kyutuynda core: 10 samples; Lake Bolshoye Toko core: 8 samples). Extraction blanks of each batch of DNA isolation were processed along with the DNA extracts. All extractions and metabarcoding PCR setup were performed in dedicated ancient-DNA laboratory facilities of the Alfred Wegener Institute, Helmholtz Centre for Polar and Marine Research, Potsdam, Germany.

#### Metabarcoding PCRs and sequencing

sedaDNA metabarcoding PCRs of CH12 extracts were performed according to the established PCR conditions described above, using primers tagged with individual eight-bp tags preceded by three variable positions (‘N’) to improve cluster formation and using 9 ng DNA; PCRs on other cores are described by von Hippel *et al*. (2021). Six technical replicates of the PCR of each extract (N = 70) and extraction blank (N = 11) were used, with one non-template control per PCR batch (N = 30), resulting in a total of 510 PCR reactions. MetaFast library preparation and paired-end sequencing at 2 x 250 bp on an Illumina MiSeq platform (Illumina, San Diego, CA, USA) was performed by a commercial service provider (Fastens SA, Geneva, Switzerland).

#### Data analyses

Data were processed and analyzed using obitools software (Boyer *et al*., 2016) as described by von Hippel *et al*. (2021). Briefly, sequences were clustered using sumaclust (Mercier *et al*., 2013) at a score threshold of 0.97 and were annotated using the obitools ecotag assignment with two databases – the latest UNITE release v. 8.2 (Abarenkov *et al*., 2010) and the EMBL Nucleotide Sequence Database (Kanz *et al*., 2005) release #142. Only reads containing both primers and both tags were kept in the dataset. The dataset was then filtered by best identity scores to count the number of unique taxa on order, family, genus levels in a range of 95% to 100% best identity. As we observed that the curves of additional numbers of taxa flattened at approximately 97%, we removed all OTUs with best identity < 97%. We retained only OTUs that a) occurred at a minimum of 10 reads per PCR replicate and b) occurred at a minimum of 100 reads across the entire dataset. Read counts were normalized by transforming each OTU’s count in a replicate to a proportion of the sum of all reads in the respective sample (proportional data per replicate). To produce proportional data per sample, reads per replicate were summed, and proportions were calculated accordingly. Shannon-Wiener diversity indices of each sample were calculated per lake on the taxonomic levels of genus, family, and order using the function ‘ddply’ of the R package plyr version 1.8.6 (Wickham, 2011).

Biases introduced through experimental procedures and/or through DNA degradation were examined in a number of ways: to assess whether we sampled the fungal diversity comprehensively and representatively, we investigated accumulation curves both for the combined PCR replicates through rarefaction in single PCRs. Specifically, to determine the number of OTUs per cumulative number of PCR replicates, we produced accumulation curves of each sample of core CH12 using the function ‘specaccum’ of the R package vegan (Oksanen *et al*., 2020). To examine whether sequencing depth was sufficient, we rarefied the data set of core CH12 using ‘rarefy’ in vegan (Oksanen *et al*., 2020) and produced rarefaction curves.

To test for replicability of results, we investigated whether dissimilarities between samples are larger than those between replicates. We used the transformed proportional data of core CH12 for a multiple response permutation procedure (MRPP) with the function mrpp (vegan; Oksanen *et al*., 2020) and Bray–Curtis dissimilarity distances.

In data produced from all lake cores, the potential effect of DNA degradation on the results was assessed both considering the GC content of recovered sequences, as well as sequence length. To assess DNA degradation-induced bias towards amplicons with higher GC content (Dabney *et al*., 2013), we examined GC proportions over time as weighted means per sample. The effect of age on weighted mean GC content was tested using a linear regression model. To determine differences in amplicon length between fungal divisions, we used an ANOVA and a Tukey’s test *post hoc*; to assess potential effects of age on amplicon length (i.e., whether longer fragments are less likely to be amplified in older samples), we fitted a linear regression of amplicon length and age, for each core. Data analyses were performed using R software, version 3.6.0 (R Development Core Team, 2019).

## Data availability statement

Raw sequencing data will be made available in an online repository upon acceptance.

## Acknowledgements

This research was funded through the 2017-2018 Belmont Forum and BiodivERsA joint call for research proposals, under the BiodivScen ERA-Net COFUND program, and with the funding organizations Deutsche Forschungsgemeinschaft (DFG grant EP-98/3-1 to L.S. Epp), Agence Nationale de la Recherche (ANR), Research Council of Norway (NFR), Formas, Academy of Finland, National Science Foundation (NSF) and the Natural Sciences and Engineering Research Council of Canada (NSERC-CRSNG). This study was supported by the ERC consolidator grant Glacial Legacy to U. Herzschuh (no. 772852). We thank the Natural History Museum of Oslo, Norway, particularly Gunnhild Marthinsen and Katriina Bendiksen, for providing vouchers for DNA extraction for the specificity tests and optimization of the primers.

## Notes

### Competing Interest Statement

The authors have declared no competing interest.

### Summary of Updates

A reference was added.

## References

Abarenkov, K., Henrik Nilsson, R., Larsson, K.-H., Alexander, I.J., Eberhardt, U., Erland, S., et al. (2010) The UNITE database for molecular identification of fungi - recent updates and future perspectives. New Phytol 186:281–285.

Adamo, M., Voyron, S., Chialva, M., Marmeisse, R., and Girlanda, M. (2020) Metabarcoding on both environmental DNA and RNA highlights differences between fungal communities sampled in different habitats. PLoS One 15:eO244682.

Baldrian, P., Větrovský, T., Lepinay, C., and Kohout, P. (2021) High-throughput sequencing view on the magnitude of global fungal diversity. Fungal Divers.

Banchi, E., Stankovic, D., Fernández-Mendoza, F., Gionechetti, F., Pallavicini, A., and Muggia, L. (2018) ITS2 metabarcoding analysis complements lichen mycobiome diversity data. Mycol Prog 17:1049–1066.

Bar-On, Y.M., Phillips, R., and Milo, R. (2018) The biomass distribution on Earth. Proc Natl Acad Sci 115:6506–6511.

Bellemain, E., Carlsen, T., Brochmann, C., Coissac, E., Taberlet, P., and Kauserud, H. (2010) ITS as an environmental DNA barcode for fungi: an in silico approach reveals potential PCR biases. BMC Microbiol 10:189.

Bellemain, E., Davey, M.L., Kauserud, H., Epp, L.S., Boessenkool, S., Coissac, E., et al. (2013) Fungal palaeodiversity revealed using high-throughput metabarcoding of ancient DNA from arctic permafrost. Environ Microbiol 15:1176–1189.

Blaalid, R., Kumar, S., Nilsson, R.H., Abarenkov, K., Kirk, P.M., and Kauserud, H. (2013) ITS1 versus ITS2 as DNA metabarcodes for fungi. Mol Ecol Resour 13:218–224.

Blackwell, M. (2011) The Fungi: 1, 2, 3… 5.1 million species? Am J Bot 98:426–438.

Booth, T. (2011) Taxonomic notes on coprophilous fungi of the Arctic: Churchill, Resolute Bay, and Devon Island. Can J Bot 60:1115–1125.

Botnen, S.S., Thoen, E., Eidesen, P.B., Krabberød, A.K., and Kauserud, H. (2020) Community composition of arctic root-associated fungi mirrors host plant phylogeny. FEMS Microbiol Ecol 96:.

Boyer, F., Mercier, C., Bonin, A., Le Bras, Y., Taberlet, P., and Coissac, E. (2016) obitools: a unix-inspired software package for DNA metabarcoding. Mol Ecol Resour 16:176–182.

Brundrett, M. (2004) Diversity and classification of mycorrhizal associations. Biol Rev Camb Philos Soc 79:473–495.

Capo, E., Giguet-covex, C., Rouillard, A., Nota, K., and Peter, D. (2020) Lake sedimentary DNA research on past terrestrial and aquatic biodiversity⍰: Overview and recommendations.

Chepstow-Lusty, A.J., Frogley, M.R., and Baker, A.S. (2019) Comparison of Sporormiella dung fungal spores and oribatid mites as indicators of large herbivore presence: evidence from the Cuzco region of Peru. J Archaeol Sci 102:61–70.

Clemmensen, K.E., Bahr, A., Ovaskainen, O., Dahlberg, A., Ekblad, A., Wallander, H., et al. (2013) Roots and Associated Fungi Drive Long-Term Carbon Sequestration in Boreal Forest. Science (80-) 339:1615–1618.

Dabney, J., Meyer, M., and Paabo, S. (2013) Ancient DNA Damage. Cold Spring Harb Perspect Biol 5:a012567–a012567.

Davis, O.K. and Shafer, D.S. (2006) Sporormiella fungal spores, a palynological means of detecting herbivore density. Palaeogeogr Palaeoclimatol Palaeoecol 237:40–50.

Deacon, J.W., Donaldson, S.J., and Last, F.T. (1983) Sequences and interactions of mycorrhizal fungi on birch. Plant Soil 71:257–262.

Deiner, K., Bik, H.M., Mächler, E., Seymour, M., Lacoursière-Roussel, A., Altermatt, F., et al. (2017) Environmental DNA metabarcoding: Transforming how we survey animal and plant communities. Mol Ecol 26:5872–5895.

Domaizon, I., Winegardner, A., Capo, E., Gauthier, J., and Gregory-Eaves, I. (2017) DNA-based methods in paleolimnology: new opportunities for investigating long-term dynamics of lacustrine biodiversity. J Paleolimnol 58:1–21.

Domsch, K.H., Gams, W., and Anderson, T.H. (2007) Compendium of Soil Fungi, second edi. Eching: IHW-Verlag.

Epp, L.S., Boessenkool, S., Bellemain, E., Haile, J., Esposito, A., Riaz, T., et al. (2012) New environmental metabarcodes for analysing soil DNA: potential for studying past and present ecosystems. Mol Ecol 21:1821–1833.

Feranec, R.S., Miller, N.G., Lothrop, J.C., and Graham, R.W. (2011) The Sporormiella proxy and end-Pleistocene megafaunal extinction: A perspective. Quat Int 245:333–338.

Ficetola, G., Coissac, E., Zundel, S., Riaz, T., Shehzad, W., Bessière, J., et al. (2010) An In silico approach for the evaluation of DNA barcodes. BMC Genomics 11:434.

Finlay, R.D. (2008) Ecological aspects of mycorrhizal symbiosis: with special emphasis on the functional diversity of interactions involving the extraradical mycelium. J Exp Bot 59:1115–1126.

Gardes, M. and Bruns, T.D. (1993) ITS primers with enhanced specificity for basidiomycetes - application to the identification of mycorrhizae and rusts. Mol Ecol 2:113–118.

Grau, O., Geml, J., Pérez-Haase, A., Ninot, J.M., Semenova-Nelsen, T.A., and Peñuelas, J. (2017) Abrupt changes in the composition and function of fungal communities along an environmental gradient in the high Arctic. Mol Ecol 26:4798–4810.

Haile, J., Froese, D.G., MacPhee, R.D.E.E., Roberts, R.G., Arnold, L.J., Reyes, A.V., et al. (2009) Ancient DNA reveals late survival of mammoth and horse in interior Alaska. Proc Natl Acad Sci 106:22352–22357.

Hawksworth, D.L. and Lücking, R. (2017) Fungal Diversity Revisited: 2.2 to 3.8 Million Species. Microbiol Spectr 5:.

Heeger, F., Bourne, E.C., Baschien, C., Yurkov, A., Bunk, B., Spröer, C., et al. (2018) Long-read DNA metabarcoding of ribosomal RNA in the analysis of fungi from aquatic environments. Mol Ecol Resour 18:1500–1514.

van der Heijden, M.G.A., Klironomos, J.N., Ursic, M., Moutoglis, P., Streitwolf-Engel, R., Boller, T., et al. (1998) Mycorrhizal fungal diversity determines plant biodiversity, ecosystem variability and productivity. Nature 396:69–72.

von Hippel, B., Stoof-Leichsenring, K.R., Schulte, L., Seeber, P.A., Epp, L.S., Biskaborn, B.K., et al. (2021) Long-term fungus–plant co-variation from multi-site sedimentary ancient DNA metabarcoding in Siberia. bioRxiv. https://doi.org/10.1101/2021.11.05.465756

James, T.Y., Stajich, J.E., Hittinger, C.T., and Rokas, A. (2020) Toward a Fully Resolved Fungal Tree of Life. Annu Rev Microbiol 74:291–313.

Kanz, C., Aldebert, P., Althorpe, N., Baker, W., Baldwin, A., Bates, K., et al. (2005) The EMBL nucleotide sequence database. Nucleic Acids Res 33:D29.

Khomich, M., Davey, M.L., Kauserud, H., Rasconi, S., and Andersen, T. (2017) Fungal communities in Scandinavian lakes along a longitudinal gradient. Fungal Ecol 27:36–46.

Klemm, J., Herzschuh, U., and Pestryakova, L.A. (2016) Vegetation, climate and lake changes over the last 7000 years at the boreal treeline in north-central Siberia. Quat Sci Rev 147:422–434.

Kumpula, J. (2001) Winter grazing of reindeer in woodland lichen pasture. Small Rumin Res 39:121–130.

Lammers, Y., Heintzman, P.D., and Alsos, I.G. (2021) Environmental palaeogenomic reconstruction of an Ice Age algal population. Commun Biol A: 220.

Lücking, R., Aime, M.C., Robbertse, B., Miller, A.N., Ariyawansa, H.A., Aoki, T., et al. (2020) Unambiguous identification of fungi: Where do we stand and how accurate and precise is fungal DNA barcoding? IMA Fungus 11:.

Lydolph, M.C., Jacobsen, J., Arctander, P., Gilbert, M.T.P., Gilichinsky, D.A., Hansen, A.J., et al. (2005) Beringian paleoecology inferred from permafrost-preserved fungal DNA. Appl Environ Microbiol 71:1012–1017.

Martin, F. (2014) The Ecological Genomics of Fungi - Wiley Online Library.

Mercier, C., Boyer, F., Bonin, A., and Coissac, E. (2013) SUMATRA and SUMACLUST: fast and exact comparison and clustering of sequences. In SeqBio 2013 workshop. pp. 27–29.

Naranjo-Ortiz, M.A. and Gabaldón, T. (2019) Fungal evolution: diversity, taxonomy and phylogeny of the Fungi. Biol Rev 94:2101–2137.

Nilsson, R.H., Anslan, S., Bahram, M., Wurzbacher, C., Baldrian, P., and Tedersoo, L. Mycobiome diversity: high-throughput sequencing and identification of fungi Nature reviews | Microbiology. Nat Rev Microbiol.

Oksanen, A.J., Blanchet, F.G., Friendly, M., Kindt, R., Legendre, P., Mcglinn, D., et al. (2020) Community Ecology Package vegan,. 2.5-7.

Parducci, L., Bennett, K.D., Ficetola, G.F., Alsos, I.G., Suyama, Y., Wood, J.R., and Pedersen, M.W. (2017) Ancient plant DNA in lake sediments. New Phytol 214:924–942.

Pedersen, M.W., Ruter, A., Schweger, C., Friebe, H., Staff, R.A., Kjeldsen, K.K., et al. (2016) Postglacial viability and colonization in North America’s ice-free corridor. Nature 537:45–49.

Powell, J.R. and Rillig, M.C. (2018) Biodiversity of arbuscular mycorrhizal fungi and ecosystem function. New Phytol 220:1059–1075.

R Development Core Team (2019) A language and environment for statistical computing, R Foundation for Statistical Computing (ed) Vienna, Austria: R Foundation for Statistical Computing.

Raper, D. and Bush, M. (2009) A test of Sporormiella representation as a predictor of megaherbivore presence and abundance. Quat Res 71:490–496.

Riaz, T., Shehzad, W., Viari, A., Pompanon, F., Taberlet, P., and Coissac, E. (2011) ecoPrimers: inference of new DNA barcode markers from whole genome sequence analysis. Nucleic Acids Res 39:e145–e145.

Ruppert, K.M., Kline, R.J., and Rahman, M.S. (2019) Past, present, and future perspectives of environmental DNA (eDNA) metabarcoding: A systematic review in methods, monitoring, and applications of global eDNA. Glob Ecol Conserv 17:e00547.

Schoch, C.L., Seifert, K.A., Huhndorf, S., Robert, V., Spouge, J.L., Levesque, C.A., et al. (2012) Nuclear ribosomal internal transcribed spacer (ITS) region as a universal DNA barcode marker for Fungi. Proc Natl Acad Sci U S A 109:6241–6.

Smith, S. and Read, D. (2008) Mycorrhizal Symbiosis, 3rd ed. Elsevier.

Spatafora, J.W., Chang, Y., Benny, G.L., Lazarus, K., Smith, M.E., Berbee, M.L., et al. (2016) A phylum-level phylogenetic classification of zygomycete fungi based on genome-scale data. Mycologia 108:1028–1046.

Stielow, J.B., Lévesque, C.A., Seifert, K.A., Meyer, W., Irinyi, L., Smits, D., et al. (2015) One fungus, which genes? Development and assessment of universal primers for potential secondary fungal DNA barcodes. Persoonia - Mol Phylogeny Evol Fungi 35:242–263.

Taberlet, P., Coissac, E., Pompanon, F., Brochmann, C., and Willerslev, E. (2012) Towards nextgeneration biodiversity assessment using DNA metabarcoding. Mol Ecol 21:2045–2050.

Talas, L., Stivrins, N., Veski, S., Tedersoo, L., and Kisand, V. (2021) Sedimentary Ancient DNA (sedaDNA) Reveals Fungal Diversity and Environmental Drivers of Community Changes throughout the Holocene in the Present Boreal Lake Lielais Svetinu (Eastern Latvia). Microorganisms 9:719.

Taylor, T.N. and Osborn, J.M. (1996) The importance of fungi in shaping the paleoecosystem. Rev Palaeobot Palynol 90:249–262.

Tedersoo, L., Anslan, S., Bahram, M., Põlme, S., Riit, T., Liiv, I., et al. (2015) Shotgun metagenomes and multiple primer pair-barcode combinations of amplicons reveal biases in metabarcoding analyses of fungi. MycoKeys 10:1–43.

Tedersoo, L. and Lindahl, B. (2016) Fungal identification biases in microbiome projects. Environ Microbiol Rep 8:774–779.

Varma, A., Prasad, R., and Tuteja, N. eds. (2017) Mycorrhiza - Function, Diversity, State of the Art, Cham: Springer International Publishing.

Wickham, H. (2011) The Split-Apply-Combine Strategy for Data Analysis. J Stat Softw 40:.

Willerslev, E., Davison, J., Moora, M., Zobel, M., Coissac, E., Edwards, M.E., et al. (2014) Fifty thousand years of Arctic vegetation and megafaunal diet. Nature 506:47–51.

Willerslev, E., Hansen, A.J., Binladen, J., Brand, T.B., Gilbert, M.T.P., Shapiro, B., et al. (2003) Diverse plant and animal genetic records from holocene and pleistocene sediments. Science (80-) 300:791–795.

Wood, J.R. and Wilmshurst, J.M. (2013) Accumulation rates or percentages? How to quantify Sporormiella and other coprophilous fungal spores to detect late Quaternary megafaunal extinction events. Quat Sci Rev 77:1–3.

