## Supplementary Fig for "Fungal biodiversity in Arctic paleoecosystems assessed by metabarcoding of lake sedimentary ancient DNA"

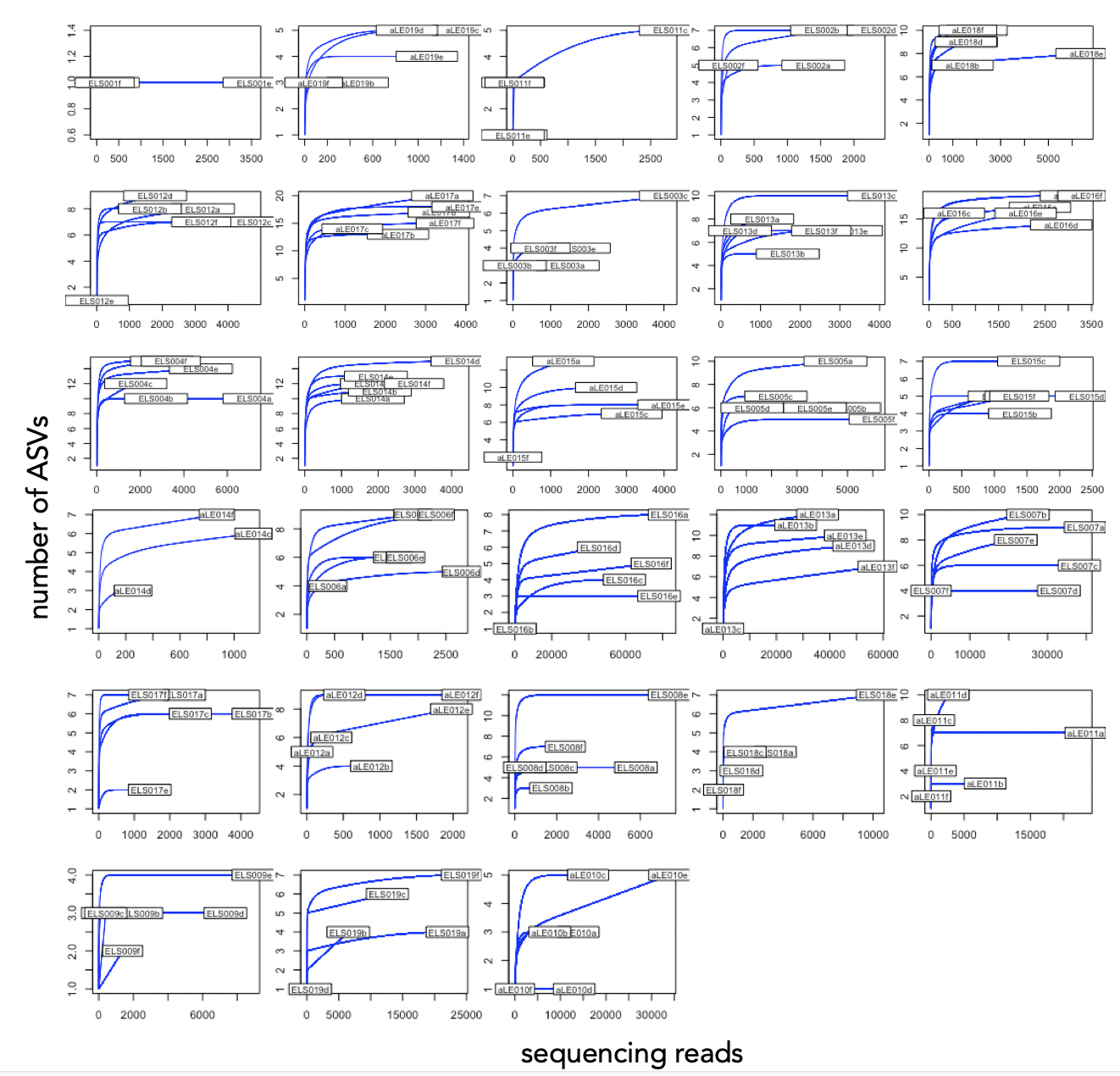

**Figure S1.** Rarefaction curves of PCR replicates of core CH12. Shown are the numbers of OTUs as a function of the number of sequencing reads.

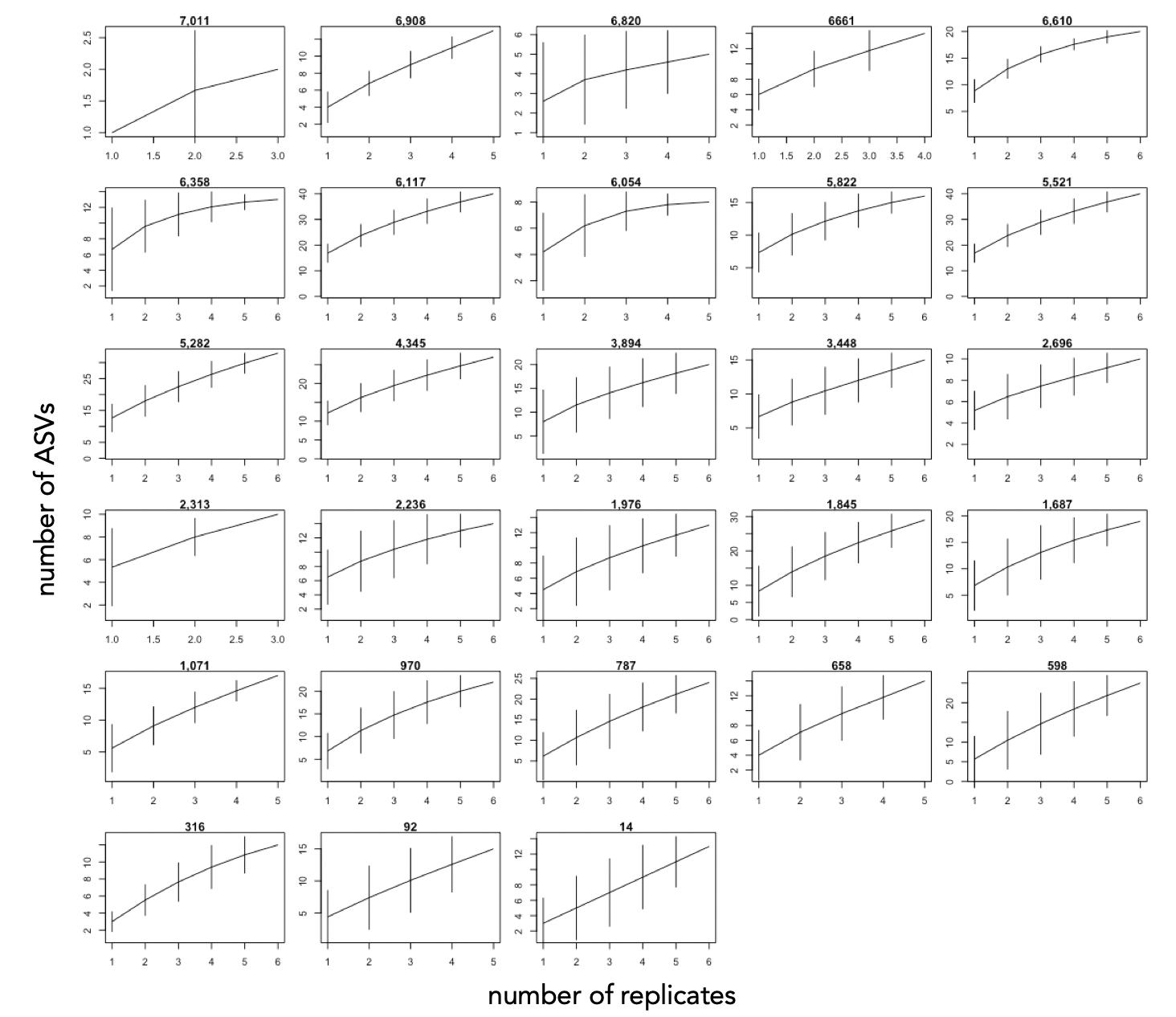

**Figure S2.** Saturation curves of number of OTUs per cumulative number of replicates in the sediment core of lake CH12. The age of each sample is indicated above the respective graph.

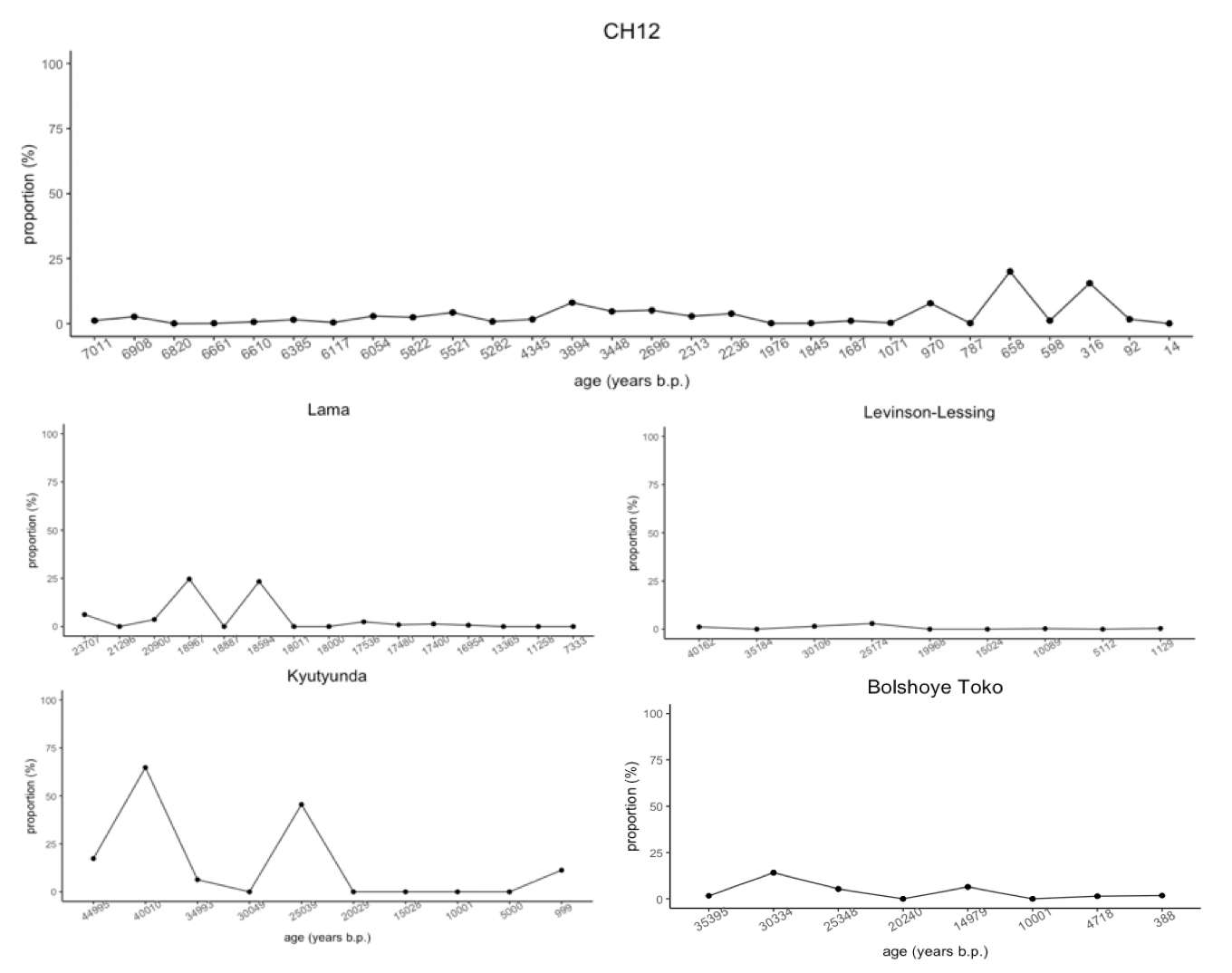

**Fig. S3.** Cumulative proportions of mold genera (*Aspergillus*, *Cladosporium*, *Mucor*, and *Penicillium*) over time in the five sediment cores.
